# GABAergic regulation of cell proliferation within the adult mouse spinal cord

**DOI:** 10.1101/2022.04.21.489053

**Authors:** Lauryn E New, Yuchio Yanagawa, Glenn A McConkey, Jim Deuchars, Susan A Deuchars

## Abstract

Manipulation of neural stem cell proliferation and differentiation in the postnatal CNS is receiving significant attention due to therapeutic potential. In the spinal cord, such manipulations may promote repair in conditions such as multiple sclerosis or spinal cord injury, but may also limit excessive cell proliferation contributing to tumours such as ependymomas. Here we show that when ambient GABA is increased in vigabatrin-treated or decreased in glutamic acid decarboxylase67-green fluorescent protein (GAD67-GFP) mice, the numbers of proliferating cells respectively decreased or increased. Thus, intrinsic spinal cord GABA levels are correlated with the extent of cell proliferation, providing important evidence for the possibility of manipulating these levels. Diazepam binding inhibitor, an endogenous protein that interacts with GABA receptors and its breakdown product, octadecaneuropeptide, which preferentially activates central benzodiazepine (CBR) sites, were highly expressed in the spinal cord, especially in ependymal cells surrounding the central canal. Furthermore, animals with reduced CBR activation via treatment with flumazenil or Ro15-4513, or with a G2F77I mutation in the CBR binding site had greater numbers of Ethynyl-2’-deoxyuridine positive cells compared to control, which maintained their stem cell status since the proportion of newly proliferated cells becoming oligodendrocytes or astrocytes was significantly lower. Altering endogenous GABA levels or modulating GABAergic signaling through specific sites on the GABA receptors therefore influences NSC proliferation in the adult spinal cord. These findings provide a basis for further study into how GABAergic signaling could be manipulated to enable spinal cord self-regeneration and recovery or limit pathological proliferative activity.

## 1. Introduction

Influences on cell proliferation in the adult brain are under intense scrutiny, with a hope of enabling CNS repair. However, increasing recognition of regional specialization means that an approach in one CNS region is unlikely to be generally applicable. In particular, populations of cells in the spinal cord with the ability to proliferate are transcriptionally distinct from their namesakes in the brain, including oligodendrocytes (Khandker et al., 2022), astrocytes (Tsai et al., 2012) and ependymal cells (MacDonald et al., 2021). Proliferation of cells in the adult spinal cord may have positive and negative effects on spinal cord health. For example, repair of myelin damaged in multiple sclerosis or spinal cord injury will benefit from production of new oligodendrocytes (Lubetzki et al., 2020). In contrast, pathological proliferation of ependymal cells can result in tumours known as ependymomas (Saleh et al., 2022) while astrocytic proliferation can cause astrocytomas (Momin et al., 2022) or contribute to axon regrowth inhibiting glial scars formation following spinal cord injury (Yang et al., 2020). It is therefore necessary to understand mechanisms controlling proliferation in the spinal cord.

Ependymal cells (ECs) surrounding the central canal (CC) of adult spinal cord could be particularly relevant for study as they possess neural stem cell (NSC) properties (Johansson et al., 1999; Martens et al., 2002; Meletis et al., 2008; Reynolds and Weiss, 1992; Sabelström et al., 2014; Sabourin et al., 2009). In the intact cord these ECs are relatively quiescent, contributing only ~4.8% of proliferating cells (Johansson et al., 1999) but this increases dramatically upon injury impacting on the ependymal cell layer (ECL) (Barnabé-Heider et al., 2010; Hamilton et al., 2009; Li et al., 2016; Meletis et al., 2008). This increased proliferation rate is potentially beneficial, due to the varied and critically important roles of the ECL. For example, in axolotl (O’Hara et al., 1992) and zebrafish (Chang et al., 2021), species that can regenerate a completely severed spinal cord, the ECL is the first part of the cord to repair. This is consistent with ECs constituting the major part of the important spinal CSF-CNS barrier (Cousins et al., 2021), as well as being a potential source of growth factors (Del Bigio, 2010).

In brain neurogenic niches, endogenous signaling regulates the production of cells so that the demand for new cells of different lineages can be tuned to meet requirements (Alfonso et al., 2012; Daynac et al., 2013; Fabbiani et al., 2020; Miyajima et al., 2021; Morizur et al., 2018; Nguyen et al., 2003; Wang et al., 2003). One signaling mechanism is through GABA, which negatively regulates proliferative activity in the subventricular zone (SVZ) and subgranular zone (SGZ) (Bolteus and Bordey, 2004; Catavero et al., 2018; Ge et al., 2007; Nguyen et al., 2003). In the SGZ, this involves local parvalbumin interneurons that release GABA (Bao et al., 2017; Song et al., 2013), which maintains the NSCs in a state of quiescence through controlling cell cycle exit (Daynac et al., 2013; Fernando et al., 2011). GABA also modulates the ability of NSCs and immature neural cells to differentiate down the specific cell lineages and migrate to new locations (Bolteus and Bordey, 2004; Tozuka et al., 2005; Trinchero et al., 2021).

The exact mechanism of how GABA controls NSC cycle entry in the brain remained elusive until recently. Negative allosteric modulation of GABAergic neurotransmission by the endogenous ligand diazepam binding inhibitor (DBI) is an important mechanism regulating NSC cell cycle exit and the levels of this protein, and its breakdown products, therefore enable a precise equilibrium between quiescence and maintenance of proliferation (Alfonso et al., 2012; Dumitru et al., 2017).

It is currently unclear whether GABA can affect proliferation and differentiation within the mammalian spinal cord, although recent data in the zebrafish suggest that non-synaptic, ambient GABA maintains spinal NSCs in quiescence (Chang et al., 2021). Activation of GABAa receptors (GABAaR) depolarizes spinal ECs in the rat (Corns et al., 2013), progenitor cells in the turtle central canal (Reali et al., 2011) and NSCs in the zebrafish (Chang et al., 2021). Considering the important role of endozepines in modulation of GABAaR, we considered here if they may be an important mechanism for control of proliferation in the spinal cord.

We show in the adult mouse spinal cord that spinal ambient GABA levels and cell proliferation are inversely proportional. Furthermore, DBI is expressed in the adult spinal cord and cell proliferation is influenced by drugs altering binding to the CBR site, indicating that endozepinergic modulation of GABAergic signaling is an important mechanism in controlling proliferation.

## 2. Materials and Methods

### 2.1 Animals

Adult wild-type C57Bl/6 mice (WT ~6-8 weeks) and glutamic acid decarboxylase 67-green fluorescent protein (GAD67-GFP) mice (Tamamaki et al., 2003) (6-8 weeks) of either sex were used in line with the UK animals (Scientific Procedures) Act 1986 and ethical standards set out by the University of Leeds Ethical Review Committee (Project licence No. P1D97A177). Every effort was made to minimize the number of animals used and their suffering.

GAD67-GFP mice (Tamamaki et al., 2003) were originally sourced from Yuchio Yanagawa. GAD67-GFP mice harbor enhanced GFP cDNA between the GAD67 5’ flanking region and the GAD67 codon start of the GAD67 promotor (Tamamaki et al., 2003). Productive breeding results in heterozygous GAD67-GFP (GAD67^+/-^) pups with GAD67 haplodeficiency due to GFP knock in. GAD67-GFP mice are seizure free and phenotypically normal despite showing a reduction in brain GABA content, as measured by high performance liquid chromatography (HPLC) (Tamamaki et al., 2003). The use of GAD67-GFP knock-in mice provided an invaluable tool to examine how reductions in ambient GABA may affect cellular proliferation and differentiation in the spinal cord

In addition, adult (~8-12 weeks) transgenic γ2F77I (C57Bl/6, γ2F77I/F77I) mice, with the point mutation F77I in the Gabrg2 gene (Cope et al., 2004), were also used.

### 2.2 Measurement of GABA by HPLC

Brain and spinal cords for HPLC were taken from adult C57Bl/6 (*N* = 3) and GAD67-GFP (*N* = 3) mice perfused transcardially with phosphate buffer (PB). Brains and spinal cords were dissected and kept in PB on ice for ~10 minutes before HPLC.

Briefly, brain and spinal cord tissue was homogenized by sonication with 0.1 M perchloric acid (PCA) (Fisher et al., 2001). GABA standards were made using 5 mM GABA in methanol:water (80:20 w/v). The solution was filtered through a 0.22 μm membrane filter. Amino acids in the samples were derivatized with o-phthaldialdehyde. GABA concentration in brain and spinal cord samples was determined by HPLC with a UV spectrophotometric detector at 360 nm.

HPLC analysis was performed with a Dionex HPLC system consisting of a P580 Pump (Dionex, USA) and Ultimate 3000 Autosampler Column Compartment with a C18 Acclaim 150 column and an ESA Coulochem III cell. The mobile phase contained 57 mM anhydrous citric acid (Fisher Scientific, UK), 43mM sodium acetate (Dionex, USA buffer containing 0.1 mM ethylenediamine tetraacetic acid (EDTA, Sigma, USA), 1mM sodium octanesulphonate monohydrate, and 10% methanol. The pH was adjusted to 4. The mobile phase was delivered at a flow rate of 0.8 ml/min, and the column temperature was set at 40°C. GABA standards were dissolved in 0.1M PCA for chromatography. The concentration of compounds was determined and analyzed using Chromeleon software.

### 2.3 In vivo drug and EdU administration

Animals received a nightly 0.1 ml intraperitoneal (IP) injection of the thymidine analogue 5-Ethynyl-2’-deoxyuridine (EdU) (10 mM) (Invitrogen, Paisley, UK) between 19:00 and 20:00 over 4 evenings. The exception to this is animals used in figure 1 which received EdU injections at midday. In addition to EdU, animals also received 0.1 ml IP of either: vigabatrin (VGB, 200mg/kg) (*N* =3), flumazenil (5mg/kg) (*N* = 3), Ro15-4513 (0.3mg/kg), 4% TWEEN-80/3%, dimethyl sulfoxide (DMSO) in sterile saline (*N* = 3), or 0.1% TWEEN-80 containing sterile saline (*N* = 6) each evening. GAD67-GFP animals and γ2F77I animals received EdU injections only.

**Figure 1.**
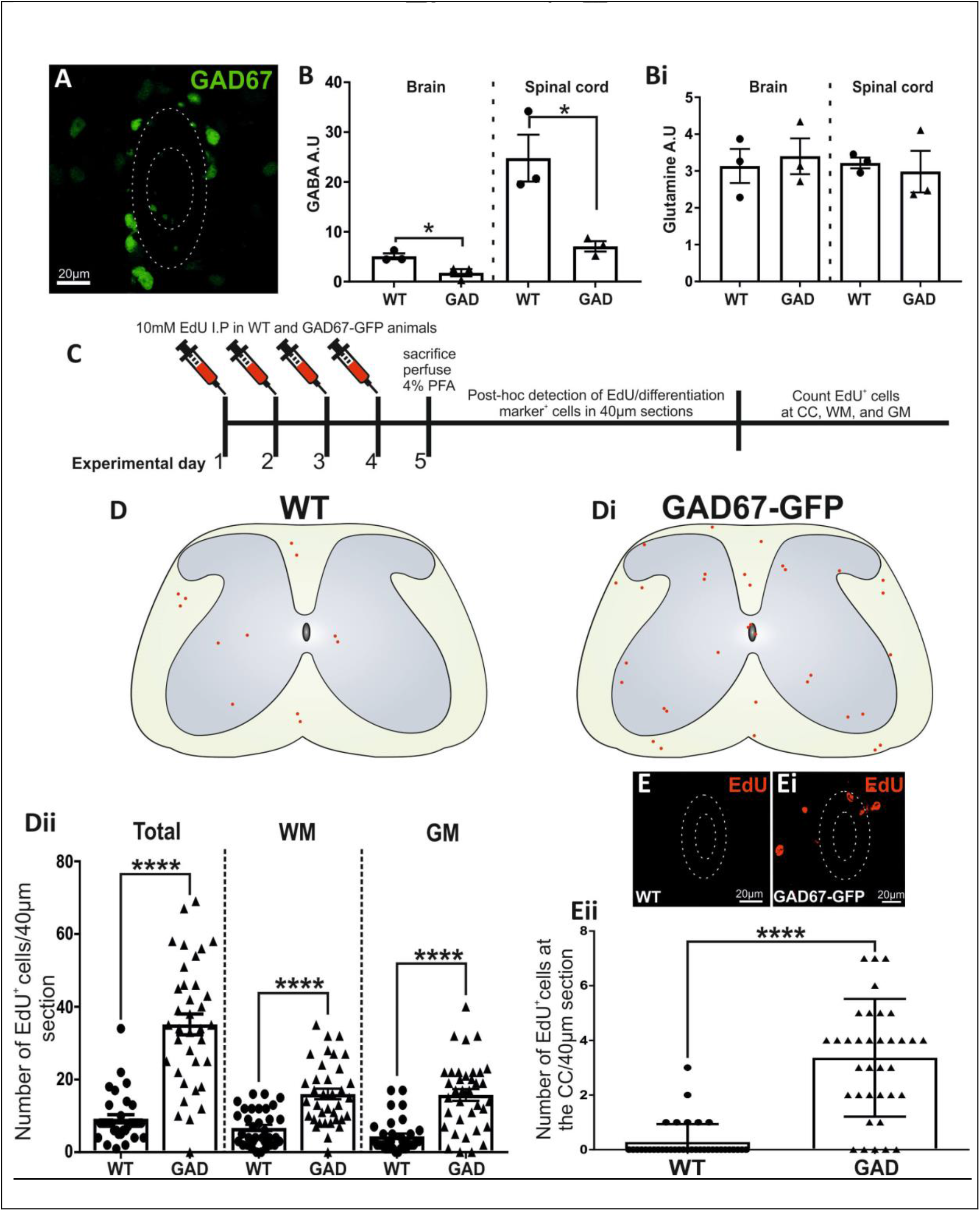
Animals with lower levels of ambient GABA in the spinal cord have increased levels of proliferation in the WM, GM, and ECL compared to WT animals. **(A):** Representative confocal image showing that GABAergic GAD67-GFP^+^ CSFcCs are present within the ECL. **(B):** HPLC analysis of GABA and **(Bi)** glutamine content in the adult spinal cord of WT vs. GAD67-GFP^+^ animals. **(C):** Schematic showing the *in vivo* experimental paradigm and post-hoc detection of proliferative EdU^+^ cells +/− differentiation markers. **(D-Di):** Representative diagrams showing the distribution of EdU-labeled cells per 40 μm spinal cord sections from **(D)** WT and **(Di)** GAD67-GFP animals. **(Dii):** Average number of total EdU^+^ cells ± SE per 40 μm section in the WM and GM combined, in the WM alone, and the GM alone in WT vs. GAD67-GFP animals. **(E):** Representative images of EdU^+^ cells at the central canal in WT and **(Ei)** GAD67-GFP animals, alongside **(Eii)** average cell counts per 40 μm section for EdU-labeled cells located in the ECL in WT vs. GADF67-GFP animals. *, *p* < 0.05, ***, *p* < 0.001, ****, *p* < 0.0001. Abbreviations: EdU, 5-ethynyl-2’-deoxyuridine; GAD67-GFP, Glutamic acid decarboxylase-67 green fluorescent protein; WT, wild type; GM: grey matter, WM; white matter; CC, central canal; PFA, paraformaldehyde; IP, intraperitoneal.

### 2.4 Tissue preparation for EdU detection and immunohistochemistry

After 5 days animals were terminally anaesthetized by IP injection of sodium pentobarbitone (60mg/kg) (Pentoject, Animalcare, UK) and perfused transcardially using 4% paraformaldehyde (PFA) as described previously (Corns et al., 2015). Spinal cords were removed and postfixed for 4-6 hours in 4% PFA at 4°C. Tissue from the thoracolumbar region (T11-L4) was sectioned at 40 μm on a vibrating microtome (Leica, VT1000S).

### 2.5 EdU localization and double labeling immunofluorescence

Labeling for EdU and differentiation markers was carried out as previously described (Corns et al., 2015). Immunofluorescence was performed with antibodies against DBI (rabbit, 1:2000, Frontier Institute, Japan), ODN (rabbit, 1:250, gift of Dr M. C. Tonon, Universitè of Rouen, France), Nestin (rat, 1:1000, BD Biosciences, Oxford, UK), CD24-FITC (mouse, 1:500, BD Biosciences), NeuN (mouse, 1:1000, Millipore, Watford, UK), Sox2, (goat, 1:1000, Santa Cruz), PanQKI, (mouse, 1:100, UC Davis/NIH Neuromab Facility, Davis, CA), S100β (rabbit, 1:750, Abcam, Cambridge, UK), glial fibrillary acidic protein (GFAP) (mouse, 1:100, Neuromab). All antibodies were diluted in PBS containing 0.1% triton (PBST) except for CD24 which was diluted in PBS. Antibodies were detected using appropriate Alexa^488^ or Alexa^555^ conjugated secondary antibodies (1:1000 in PBS) (Invitrogen). Sections were mounted on glass slides using vectashield plus DAPI (4’,6-diamidino-2-phenylindole) (Vector labs, Peterborough, UK).

### 2.6 Cell counts, image capture and processing

Cell counts, including colocalization of EdU and specific cellular markers, were performed manually at x40 magnification using a Nikon E600 microscope equipped with epifluorescence at custom settings. The numbers of EdU^+^ cells, in addition to EdU^+^ cells immunopositive for specific markers of differentiation, were counted in the white matter (WM), grey matter (GM), and CC region of every 3^rd^ 40 μm section. Investigators were blinded to the experimental group during cell counting.

Sections were imaged using a Zeiss LSM880 upright confocal laser scanning microscope with Airyscan equipped with argon and He-Ne lasers and a 40x Fluor oil objective. Images were acquired using Carl Zeiss ZEN software (Zeiss microscopy, Germany). CorelDRAW 2018 graphics suite was used for image manipulation, including correction of brightness, contrast, and intensity, and production of figures. Figures are single plane confocal images.

### 2.7 Statistical Analysis of Data

Data were collated in Microsoft Excel and analyzed using GraphPad Prism 7 (GraphPad Software, California, USA). Cell counts are given as mean number of EdU^+^ cells per 40μm section where *N* = number of animals in each experimental group and *n* = number of sections counted for each condition. Data are presented as means ± SE and for statistical analysis. Student’s t-tests and one-way ANOVA with post hoc Tukey’s test were carried out to determine significant difference in the levels of proliferation between groups, including any changes in the percentage of colocalization between EdU^+^ cells and specific cellular markers. Data were considered significant when *p* <0.05 (denoted by *); *p* <0.01 (denoted by **), *p* <0.005 (denoted by ***); or *p* <0.001 (denoted by ****).

## 3. Results

### 3.1 GAD67-GFP transgenic mice have lower levels of GABA and greater numbers of proliferative cells in the spinal cord than WT mice

HPLC analysis showed that GABA content was significantly reduced in both the brain and spinal cord of GAD67-GFP animals compared to WT C57BL/6 mice (t(4) = 3.47, *p* < 0.0256 and t(4) = 3.68, *p* < 0.0213, for brain and spinal cord, respectively, *N* = 3 animals, Fig. 1A,B). Brains of GAD67-GFP mice had on average 65% less GABA than WT, whereas spinal cords had 71% less GABA than WTs. Glutamine content, also measured by HPLC was not significantly different between groups (Fig. 1Bi).

Mapping EdU^+^ cells within the entire thoracolumbar spinal cord showed that GAD67-GFP animals had significantly greater total numbers of EdU^+^ cells compared to WT animals (35.1 ± 2.8 vs.9.2 ± 1.1, t(70) = 7.1, *p* = 8.85 × 10^-10^, *n* = 36 sections) (Fig. 1D).

In the WM alone (Fig. 1Dii) GAD67-GFP animals possessed significantly greater numbers (t(70) = 4.47, *p* = 2.97 x 10^-5^) of EdU^+^ cells (16 ± 1.1) compared to WT animals (6.7 ± 0.9 *n* = 36 sections) (Fig 1Dii). Similarly, in the GM, not including the ependymal cell layer (ECL), there were significantly greater numbers of EdU^+^ cells in GAD67-GFP animals compared to WT (15.8 ± 1.6 vs. 4.4 ± 0.8 cells, t(70) = 5.64, *p* = 3.3 x 10^-7^, *n* = 36 sections).

Proliferative EdU^+^ cells were rarely found in the ECL of WT animals (Fig. 1E), however GAD67-GFP animals had significantly greater numbers of proliferative cells localized in the ECL (3.4 ± 0.4 vs. 0.27 ± 0.1, respectively, t(70) = 8.21, *p* = 7.37 x 10^-12^, student’s t-test, *n* = 36 sections; Fig. 1Ei-ii).

These findings illustrate that GAD67-GFP animals have higher baseline levels of proliferation compared to WTs.

### 3.2 GABA transaminase inhibitor vigabatrin treated animals had significantly lower levels of proliferation compared to vehicle treatment

GAD67-GFP animals with decreased GABA levels exhibited increased proliferation and so it was hypothesized that an increase in GABA may result in decreased proliferation. However, as baseline proliferation in the intact adult spinal cord is already low, particularly at the ECL, we sought to determine whether injecting animals with EdU during the evening, when they are awake and active, would label a greater proliferative pool. This would then allow us to more easily see reductions in proliferation.

Indeed, animals injected with EdU during the dark phase of the housing light cycle, between 19:00 and 20:00, possessed greater numbers of both total EdU+ cells (75 ± 3 vs. 13.2 ± 1.9, t(88) = 15.3, *p* = 1.73 x 10^-26^, *n* = 54 and 36 sections, respectively, Fig. 2B) and EdU+ cells in the ECL (2.3 ± 0.2 vs. 0.3 ± 0.1, t(88) = 7.712, *p* = 1.3 x 10^-12^. *n* = 54 and 36 sections, respectively, Fig. 2Bi) compared to animals injected at midday. As a result of this finding, all further injections were performed between 19:00 and 20:00.

**Figure 2.**
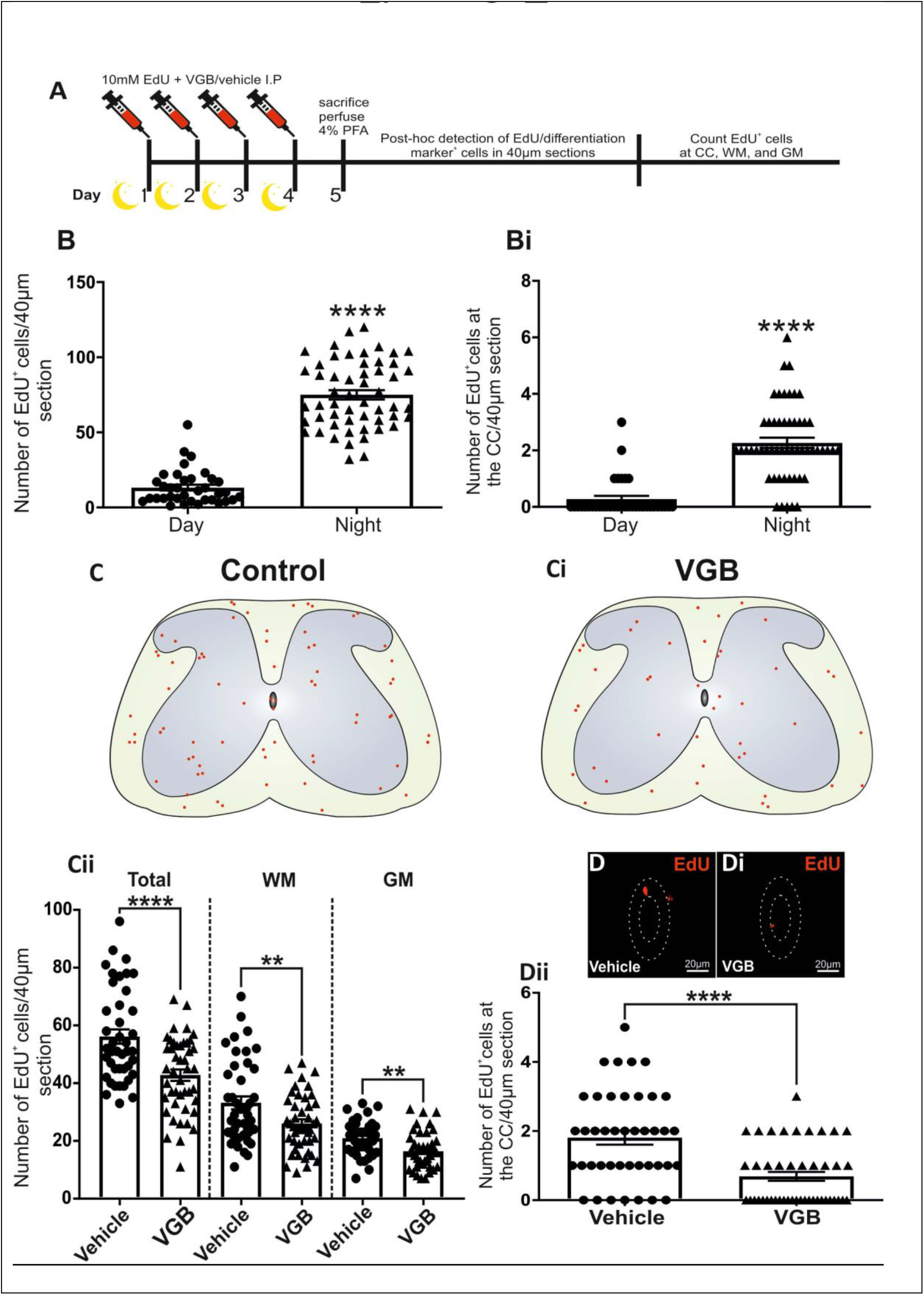
In animals treated with the GABA transaminase inhibitor VGB there were significantly lower numbers of EdU^+^ cells in the spinal cord WM, GM and ECL compared to vehicle treated controls. **(A):** Schematic showing the *in vivo* experimental paradigm and post-hoc detection of proliferative EdU^+^ cells +/− markers of differentiation. **(B-Bi):** Average number of EdU+ cells ± SE in the entire spinal cord section (comprising WM and GM) and at the CC of animals treated with EdU during daylight hours vs animals treated during dark hours **(C-Ci):** Representative diagrams showing the distribution of EdU-labeled cells in spinal cord sections from vehicle- and VGB-treated animals. **(Cii):** Average number of total EdU^+^ cells ± SE in the WM and GM combined, in the WM alone, and the GM alone in vehicle- vs. VGB-treated animals. **(D-Dii):** Representative images of EdU^+^ cells in ECL in vehicle- and VGB-treated animals, alongside average cell counts ± SE for EdU-labeled cells in ECL in vehicle vs. VGB treatment in WT vs. GADF67-GFP animals. **, *p* < 0.01, ****, *p* < 0.0001. Abbreviations: EdU, 5-ethynyl-2’-deoxyuridine; GAD67-GFP, VGB, vigabatrin; GM: grey matter, WM; white matter; CC, central canal; IP, intraperitoneal.

Animals treated with VGB, at a dose that has previously been shown to significantly enhance brain concentrations of GABA in mice (Świąder et al., 2020; Zhang et al., 2013), had significantly fewer EdU^+^ cells in total compared to vehicle treated animals (42.8 ± 2 vs. 56.2 ± 2.5, respectively, t(85) = 4.26, p = 5.33 x10^-5^, *n* = 45, Fig. 2C-Cii), exhibiting a 24% difference in proliferation compared to vehicle treatment. These differences were not confined to one specific parenchymal region as VGB treated animals exhibited significantly less proliferation in both the WM and GM (26 ± 1.5 vs.32.2 ± 2.2, t(85) = 2.73, p = 0.0077, and 16.4 ± 0.9 vs. 21 ± 0.9, t(85) = 3.56, p = 0.0006, for WM and GM, respectively, n = 45 Fig. 2Cii) compared to vehicle treated littermates.

In vehicle treated animals the average number of EdU^+^ cells in ECL per section was 1.8 ± 0.2 cells (*n* = 45 sections), which was higher (t(85) = 4.72, p = 9.23 x 10^-6^) than VGB treated animals. (0.68 ± 0.13 EdU^+^ cells, *n* = 45 sections, Fig. 2D-Dii). VGB treated animals possessed 62% fewer cells in ECL compared to vehicle treated animals.

Thus, increasing ambient GABA by VGB treatment results in a significantly lower number of proliferating cells compared to vehicle treated animals.

### 3.3 The endogenous GABAaR ligand DBI is strongly expressed in ECL of adult spinal cord

Intense DBI immunoreactivity (DBI-IR) was preferentially located in ECL (Fig. 3Ai), with less intense staining in both the WM and GM (Fig. 3A-Ai). Much like the brain, glial cells of the spinal cord also express DBI (Fig. 3E). At the dorsal and ventral poles of the CC, DBI is expressed within nestin positive radial glia (Fig. 3B-Bii).

**Figure 3.**
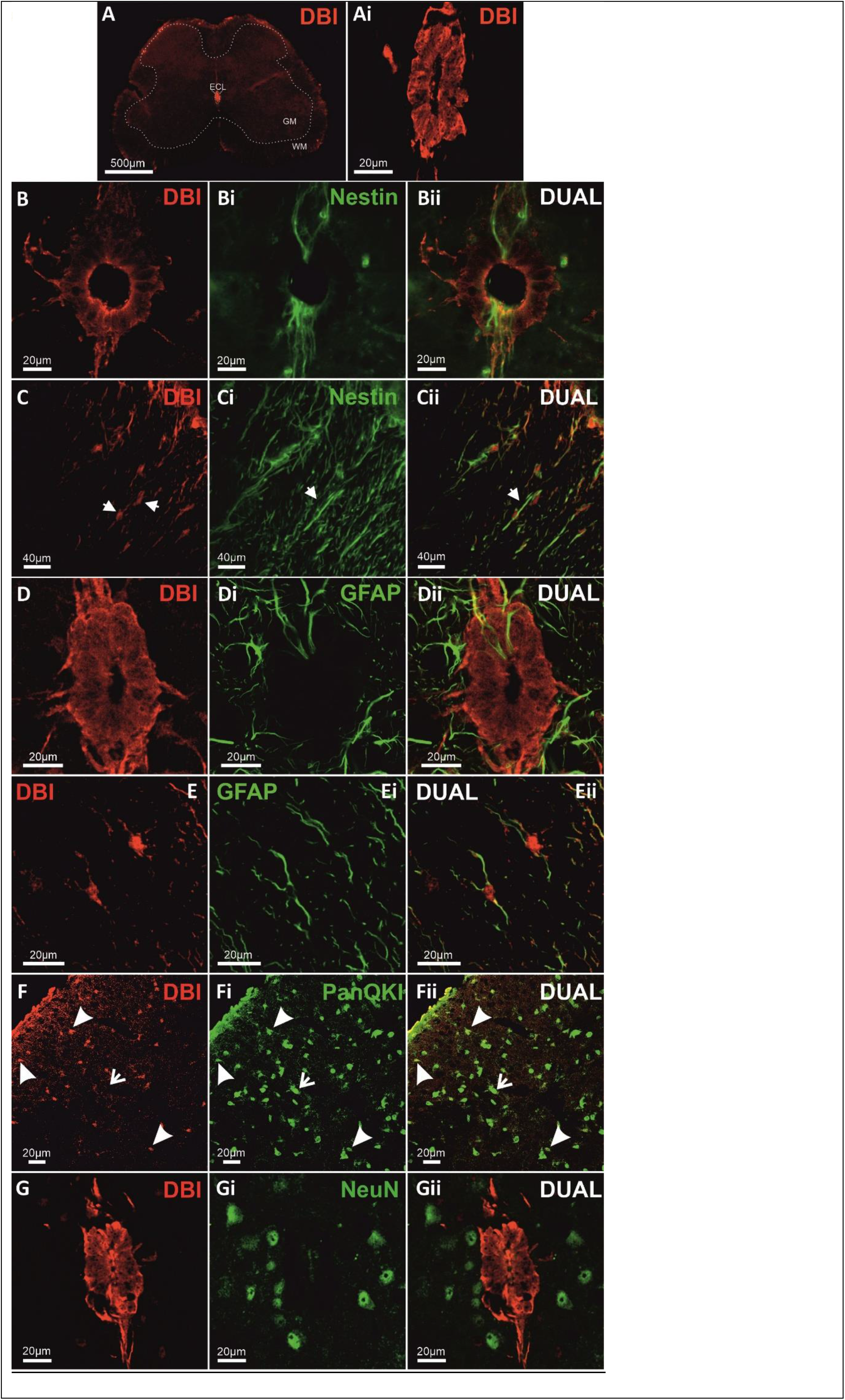
DBI is also expressed in the adult spinal cord, including radial glia, astrocytes, and oligodendrocytes but is absent from mature NeuN^+^ neurons. **(A-Ai):** Confocal images showing that DBI is expressed in the adult spinal cord, with strong immunoreactivity in ECL. Dotted lines in A denote the different spinal regions **(B-Bi):** Representative confocal images of DBI, nestin and dual labeling in ECL **(C-Cii)**: Confocal images of DBI and nestin co-labeling within radial glia of the spinal cord dorsal horn **(D-Dii):** Confocal images showing labeling for DBI in ECL and GFAP+ glial fibers within lamina X **(E-Fii):** Confocal images showing DBI+ cells are colocalized with markers of astrocytes (GFAP) and oligodendrocytes (PanQKI) in WM. **(G-Gii):** Confocal images showing that DBI is not expressed in NeuN+ mature neurons in lamina X. Closed arrows denote colocalized cells, open arrows denote non-colocalized cells. Abbreviations: DBI, diazepam binding inhibitor; GFAP, glial fibrillary acidic protein; NeuN, neuronal nuclei.

In the WM, colocalization of DBI and nestin was found in cell bodies and fibers of nestin^+^ astrocytes (Fig. 3C-Cii). GFAP^+^ astrocytic fibers in lamina X were in close apposition to the DBI-IR ECL (Fig. 3D-Dii). GFAP^+^ fibers also often passed through to contact the CC lumen. In contrast to the lack of DBI and GFAP colocalization in the ECL, in the WM and GM DBI was present within GFAP^+^ astrocytes (Fig. 3E-Eii). Double labeling for DBI and the oligodendrocyte marker PanQKI shows that DBI is also present in some WM oligodendrocytes (Fig. 3F-Fii).

DBI was not found in mature neurons of lamina X (Fig. 3G-Gii), nor in any other population of mature NeuN+ neurons in any other region of the spinal cord (data not shown).

### 3.4 DBI is expressed within ependymal cells of the CC neurogenic niche

An antibody against the cell adhesion molecule CD24, which labels ECs and cerebrospinal fluid contacting cells (CSFcCs), was used alongside DBI (Fig. 4A-Aii). Results illustrated that DBI was preferentially expressed in ECs. Areas which are CD24^+^/DBI^-^ are likely to represent CSFcC cell bodies present in the EC layer (Fig. 4A-Aii). Using GAD67-GFP mouse tissue to visualize GFP+ GABAergic CSFcCs alongside DBI confirmed that GFP^+^ CSFcC cell bodies are DBI^-^ (Fig. 4B-Bii). DBI was also absent from CSFcC bulbs within the CC lumen (Fig. 4B-Bii). Sagittal sections of the ECL improved visualization of the CC lumen and confirmed that DBI was not expressed in either CSFcC cell bodies or end bulbs along the entire spinal cord rostrocaudal axis (Fig. 4C-Cii).

**Figure 4.**
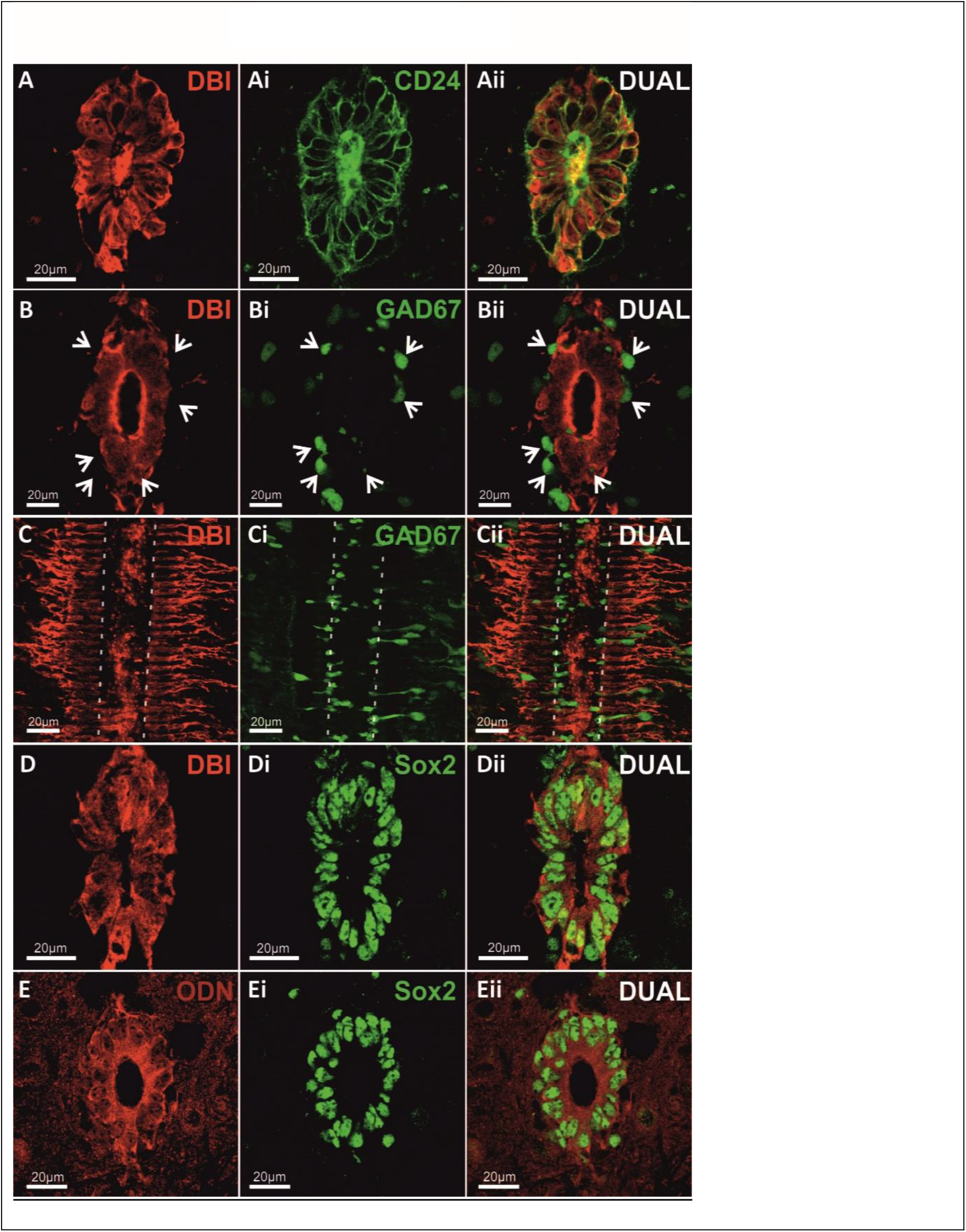
DBI-immunoreactivity is found in Sox2^+^ ependymal cells surrounding the spinal cord central canal but is absent from GAD67-GFP^+^ CSFcCs. **(A-Aii)**: Representative confocal images showing DBI^+^ cells, CD24^+^ ependymal cells and CSFcCs and dual labeling for both in ECL. **(B-Cii)**: Confocal images of DBI^+^, GAD67-GFP^+^ CSFcCs, and dual labeling in ECL, in both **(B-Bii)** transverse and **(C-Cii)** sagittal planes. Dotted outline denotes CC lumen. Open arrows denote non-colocalized cells. **(D-Eii):** Confocal images of DBI and the DBI-derived fragment ODN and Sox2^+^ stem cells in ECL. Abbreviations: DBI, diazepam binding inhibitor; CD24, cluster of differentiation 24; GAD67, glutamic acid decarboxylase-67; ODN, octadecaneuropeptide.

Sox2^+^ nuclei of ECs are located in the cytoplasm of DBI-IR cells (Fig. 4D-Dii). The full length DBI pro-peptide is a precursor to octadecaneuropeptide (ODN), a smaller fragment of DBI which also has high affinity for GABAaR/CBR (Slobodyansky et al., 1989) and replicates the effects of DBI at CBR (Alfonso et al., 2012; Dumitru et al., 2017). There was also colocalization of ODN with Sox2^+^ ECs (Fig. 4E-Eii). Furthermore, whilst some CSFcCs are also Sox2^+^, these are easily distinguished from ECs due to the absence of DBI and ODN, and their less intense labeling (Fig. 4D-Eii). Thus, DBI-IR is confined to the ECs and absent from CSFcCs.

### 3.5 Modulation of the GABAaR CBR site affects the numbers of EdU+ cells in the spinal cord

Three experimental conditions were used to assess the effects of manipulation of the CBR site, and therefore GABAaR function, upon baseline proliferation in the intact adult spinal cord. 1. Treatment with the CBR site competitive antagonist flumazenil vs vehicle treatment, 2. Treatment with the CBR site inverse agonist Ro15-4513 vs. vehicle treatment, or 3. Transgenic mutagenesis to reduce ligand binding affinity at CBR in G2F77I mice vs. WT littermates.

Animals treated with the CBR specific antagonist flumazenil (Fig. 5A) exhibited significantly higher total numbers of EdU^+^ cells within the spinal cord compared to vehicle treated animals (132.1 ± 4.4 vs. 112 ± 2.1 respectively, t(86) = 4.06, *p* = 0.00011, *N* = 3 *n* = 45, Fig. 5B). Flumazenil treated mice had 18% more EdU^+^ cells compared to vehicle. These effects were limited to proliferating cells within the WM (77.3 ± 3.6 vs. 63 ± 1.7 cells, for flumazenil and vehicle treated animals, respectively, t(86) = 3.53, *p* = 0.0007, *n* = 45, Fig. 5B) as increases in EdU^+^ cells within the GM did not reach statistical significance. Flumazenil treated animals also exhibited significantly greater levels of proliferation (5.7 ± 0.4 EdU+ cells) in ECL compared to vehicle, (2.5 ± 0.2 EdU^+^ cells) (t(86) = 7.54, *p* = 4.4 x 10^-11^, *n* = 45), a change of 128% (Fig. 5B-Bii).

**Figure 5.**
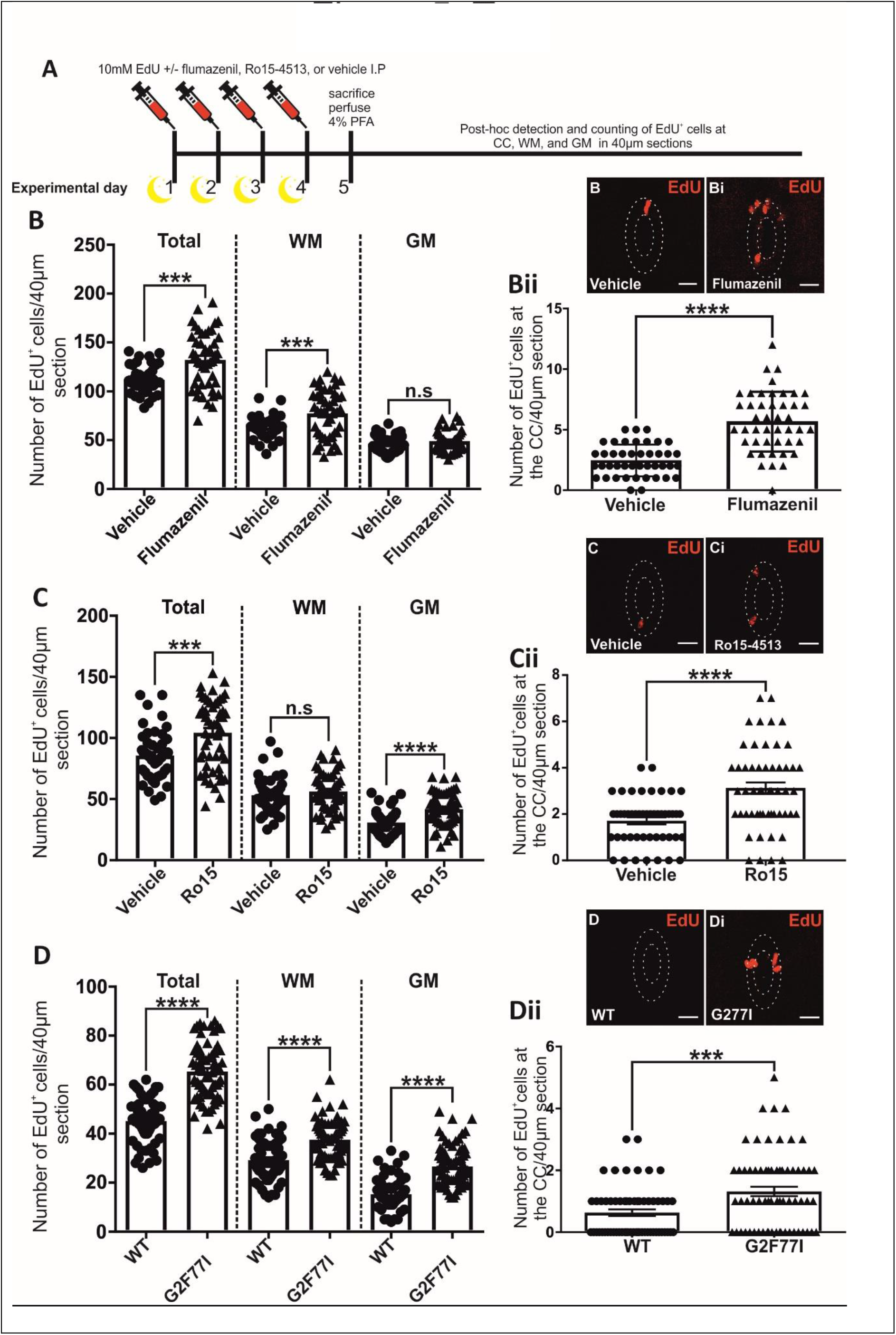
Animals with either pharmacological or transgenic modulation of the CBR site of GABAaR exhibited higher numbers of EdU-labeled cells. **(A):** Schematic showing the *in vivo* experimental paradigm and post-hoc detection of proliferative EdU^+^ cells. **(B):** Average number of total EdU^+^ cells ± SE in the WM and GM combined, in the WM alone, and the GM alone per 40 μm section in vehicle- vs. flumazenil-treated animals. **(Bi-Bii):** Representative confocal images of EdU+ cells in the ECL and scatter plots of average EdU^+^ cell counts ± SE per 40μm section in ECL following vehicle vs. VGB treatment. **(C):** Average number of total EdU^+^ cells ± SE in the WM and GM combined, in the WM alone, and the GM alone per 40 μm section in vehicle- vs. Ro15-4513-treated animals. **(Ci-Cii):** Representative confocal images of EdU+ cells in the ECL and scatter plots of average EdU^+^ cell counts ± SE per 40 μm section in ECL following vehicle vs. Ro15-4513 treatment. **(D):** Average number of total EdU^+^ cells ± SE in the WM and GM combined, in the WM alone, and the GM alone per 40 μm section in WT vs. G2F77I animals. **(Di-Dii):** Representative confocal images of EdU+ cells in the ECL and scatter plots of average cell counts of EdU^+^ cells present in ECL per 40μm section in WT animals compared to G2F77I animals. n.s, *p* > 0.05, ***, *p* < 0.001, ****, *p* < 0.0001. Abbreviations: EdU, 5-ethynyl-2’-deoxyuridine; CC, central canal; WM, white matter; GM, grey matter; Ro15, Ro15-4513; WT, wild type; IP, intraperitoneal.

Those animals treated with Ro15-4513, a weak partial inverse agonist at CBR, had significantly higher numbers of EdU^+^ cells in the whole spinal cord compared to vehicle treated animals (104.2 ± 3.8 vs. 85.6 ± 2.9 cells for Ro15-4513 and vehicle treated mice, respectively, t(100) = 3.84, *p* = 0.0002, *n* = 54, Fig. 5C-Cii). These results were similar to those in flumazenil treated animals, however there were also significantly more EdU^+^ cells in the GM following Ro15-4513 treatment compared to vehicle (41.8 ± 1.8 vs. 30.7 ± 1.4 cells respectively, t(100) = 4.71, *p* = 8 x 10^-6^, *n* = 54). Ro15-4513 induced a 22% higher number in total EdU^+^ cells and a 36% increase in the GM specifically compared to vehicle (Fig. 5C). Animals treated with Ro15-4513 also had significantly more EdU^+^ cells in ECL (3.1 ± 0.2 cells vs. 1.7 ± 0.2 cells, t(100) = 4.88, *p* = 4.1 x 10^-6^, *n* = 54). Treatment with Ro15-4513 therefore resulted in 82% more proliferative cells in ECL compared to vehicle treated animals (Fig. 5C-Cii).

Similarly, G2F77I animals, in which the mutation in the CBR binding site prevents endozepine binding, exhibited significantly greater average numbers of total EdU^+^ cells per section compared to control animals (65.9 ± 1.4 cells vs. 45.1 ± 1.1 cells in G2F77I mice and WT mice, respectively, t(124) = 11.1, *p* = 2.37 x 10^-20^, *n* = 60, Fig.5D). G2F77I mutant mice also had significantly higher numbers of EdU^+^ cells in both the WM and the GM (37.5 ± 0.9, *p* = 2.22 x 10^-8^ and 26.6 ± 1, t(130) = 5.96, *p* = 7.51 x 10^-15^ for WM and GM respectively) compared to WT controls (29.2 ± 1.1, and 15.3 ± 0.8 for WM and GM, respectively, Fig. 5D).

Proliferation in ECL was also significantly greater in G2F77I mice than in WT control animals (1.3 ± 0.2 EdU^+^ cells in ECL in G2F77I mice vs. 0.63 ± 0.1 EdU^+^ cells in ECL in WT animals (*n* = 60, t(124) = 3.64, *p* = 0.0004) (Fig. 5D-Dii)).

Thus, both pharmacological and transgenic manipulation of binding to the CBR site within GABAaR increases proliferation within the intact adult spinal cord.

### 3.6 The proportion of newly proliferated cells going down the glial cell lineage was lower following CBR modulation

In all conditions PanQKI^+^/EdU^+^cells were frequently observed in WM and GM (Fig. 6A-Aii), accounting for the greatest population of newly proliferated cells. Between 31-36.7% of newly proliferated EdU^+^ cells also expressed the oligodendrocyte marker PanQKI (Fig. 6B-Bii). Ro15-4513 treated animals showed a non-significant decrease in the percentage of PanQKI^+^/EdU^+^in the whole cord compared to vehicle treated animals (32.34 ± 1.7% vs. 36.67 ± 1.8%, t(16) = 1.7, *p* = 0.1, *n* = 9) Fig. 6B). G2F77I mice however had a significantly smaller proportion of PanQKI^+^/EdU^+^cells than WT mice (25.5 ± 2.2% vs. 33.7 ±1.7%, t(25) = 2.79, *p* = 0.01, *n* = 12, Fig. 6Bi). There was also a significantly smaller percentage of PanQKI^+^/EdU^+^cells in the GM of G2F77I mice compared to that of WT littermates (19.1 ± 1.8% vs. 29.9 ± 4.1%, t(25) = 2.6, *p* = 0.015, *n* = 12, Fig. 6Bi).

**Figure 6.**
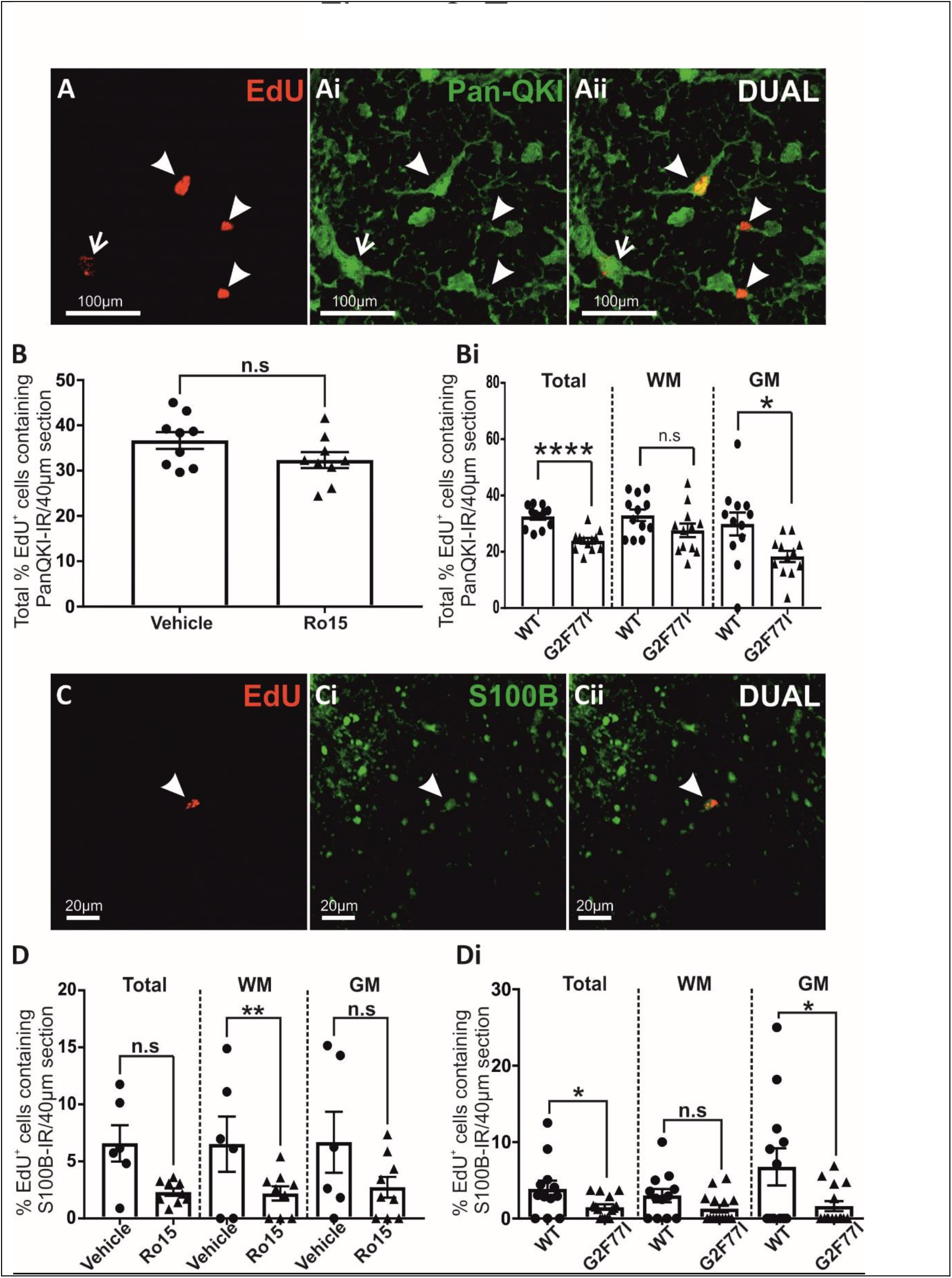
CBR modulation alters oligodendrocyte and glial cell differentiation in the adult spinal cord. **(A-Aii):** Representative confocal images of colocalization of EdU+ and Pan-QKI in the dorsal WM of the spinal cord. Closed arrows denote colocalized cells, open arrows denote non-colocalized cells. **(B):** Average percentage ± SE of EdU and PanQKI co-labeled cells per 40μm section in the WM and GM combined in vehicle vs. Ro15-4513 treated animals. **(Bi)**: Average percentage ± SE of EdU and PanQKI co-labeled cells per 40μm section in the WM and GM combined, GM alone and WM alone in WT and γ2F77I animals. **(C-Cii):** Representative confocal images of colocalization of EdU+ and S-100β in the spinal cord. Closed arrow denotes colocalized cells. **(D)**: Average percentage ± SE of EdU and S-100β co-labeled cells per 40μm section in the WM and GM combined, GM alone and WM alone in vehicle vs. Ro15-4513 treated animals. **(Di)**: Average percentage ± SE of EdU and S-100β co-labeled cells per 40μm section in the WM and GM combined, GM alone and WM alone in WT and γ2F77I animals. n.s - non-significant, *p* > 0.05, * *p* < 0.05, **, *p* < 0.01, ****, *p* < 0.0001. Abbreviations: EdU, 5-ethynyl-2’-deoxyuridine; WM, white matter; GM, grey matter; Ro15, Ro15-4513; WT, wild type.

There were also EdU^+^ cells which were also S100β^+^ (Fig. 6C-Cii), however this was a smaller percentage than that of PanQKI^+^/EdU^+^cells. Animals treated with Ro15-4513 showed a reduction in the percentage of S100β^+^/EdU cells in whole spinal cord compared to vehicle treated animals (2.4 ± 0.4% vs. 5 ± 1%, t(13) = 2.82, *p* = 0.014, *n* = 9, Fig. 6D). In G2F77I mice the proportion of total EdU^+^ cells which were co-labeled with S100β was also decreased (1.5 ± 0.4% vs. 3.9 ± 1.1% for G2F77I and WT animals, respectively, t(25) = 2.23, *p* = 0.03, *n* = 12), this change was specific to the GM (1.6 ± 0.7% vs. 6.8 ± 2.4%, t(25) = 2.25, *p* = 0.0333, *n* = 12, Fig. 6Di).

These results indicate that whilst both G2F77I and Ro15-4513 treated animals exhibit significantly greater levels of proliferation in the adult spinal cord compared to vehicle, the differentiation of these cells down either oligodendrocytic or astrocytic lineage was lower compared to WT animals.

### 3.7 Ro15-4513 treated animals had higher numbers of Sox2+ NSCs in the spinal cord

Sox2^+^/EdU^+^ cells were the second largest population of new EdU+ cells, indicating that a substantial proportion of the EdU^+^ cells in both control and experimental groups remained as undifferentiated NSCs expressing the stem cell marker Sox2. EdU^+^ cells in the ECL were consistently immunoreactive for Sox2, confirming the stem cell identity of these newly proliferated cells within the ECL (Fig. 7A).

**Figure 7.**
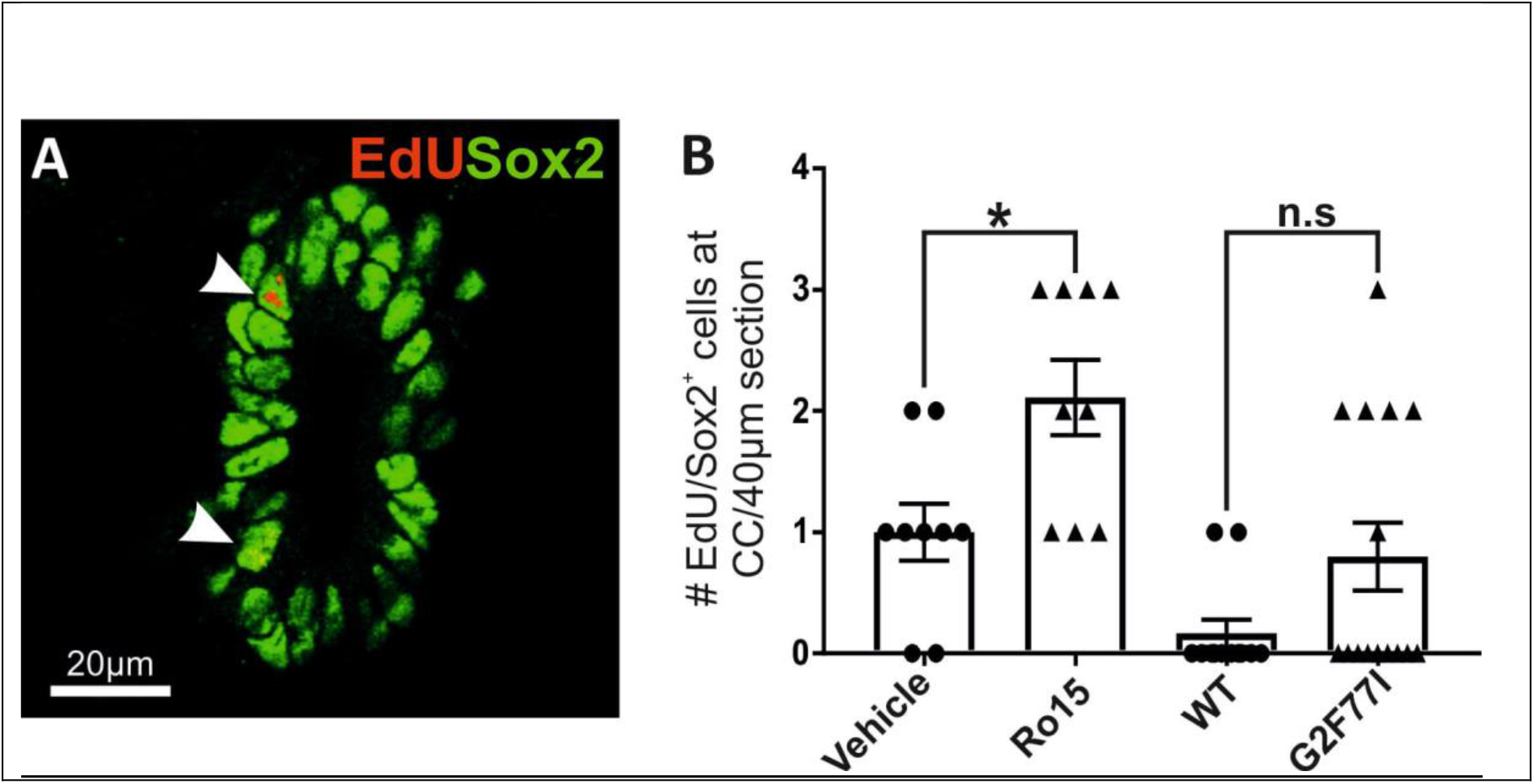
Animals treated with the CBR modulator Ro15-4513 possessed more newly proliferated cells which remained in a stem cell state compared to control animals. **(A):** Confocal image of colocalization of EdU+ and Sox2+ cells within the ECL. Closed arrows denote colocalized cells. **(B):** Average number of EdU/Sox2+ cells in the ECL in Ro15-4513 treated and G2F77I animals compared to control. There were no significant differences in numbers of double labeled cells in G2F77I mice compared to WT. n.s, p > 0.05, * p < 0.05. Abbreviations: EdU, 5-ethynyl-2’-deoxyuridine; CC, central canal; WM, white matter; GM, grey matter; Ro15, Ro15-4513

Flumazenil treated animals did not show any significant differences in the percentage of EdU^+^ cells which were also Sox2^+^ in either the WM, GM, or ECL, compared to vehicle treated animals (data not shown). The total numbers of EdU^+^ cells still residing in an undifferentiated state of stem cell readiness were also not significantly different between WT and G2F77I animals (Fig. 7B).

Ro15-4513 treated animals however exhibited an increase in the percentage of Sox2^+^/EdU^+^cells in the GM compared to vehicle treated animals. Animals treated with Ro15-4513 also had significantly more Sox2^+^/EdU^+^cells in ECL compared to vehicle treated animals (2.2 ± 0.4 cells vs. 1.1 ± 0.3 cells, *p* = 0.04, t(16) = 2.18, *n* = 9, Fig. 7B). Ro15-4513 treated animals therefore possessed more newly proliferated cells which remained in an undifferentiated stem-like state in both the GM, and the neurogenic niche of the CC.

## 4. Discussion

Findings from this study indicate that GABA is an important modulator of cell proliferation within the adult spinal cord, and that the central benzodiazepine receptors (the binding sites of both DBI and ODN) are tonically activated to dampen endogenous proliferation. Whilst previous work demonstrated that ECs exhibit GABAergic responses (Corns et al., 2013; Chang et al., 2021), results discussed here show that ambient GABA, in concert with endozepines, can modulate proliferation in the adult spinal cord within and beyond the ECL.

We show that the levels of ambient GABA and spinal cord proliferation are inversely proportional to one another, suggesting that cell proliferation is regulated by similar niche signalling mechanisms to brain where inhibitory GABAergic tone suppresses proliferation. Although ECs possess NSC properties they are relatively quiescent, raising the question as to whether high ambient GABA in the CC niche may contribute to EC quiescence and thus manipulation of GABAergic signalling is an exquisitely simple way to control proliferative activity.

### 4.1 How can GABA influence EC proliferation?

Given the important functions of ECs outlined in the introduction, including that a subset are the potential neural stem cells of the CC neurogenic niche (Barnabé-Heider et al., 2010; Johansson et al., 1999; Stenudd et al., 2022), it is important to consider how GABA is influencing their proliferation. Within other neurogenic niches, GABA excites a number of progenitor cells, including those in the ventricular zone of the developing rat brain (LoTurco et al., 1995), the mammalian postnatal SVZ (Liu et al., 2005) the mammalian postnatal SGZ (Tozuka et al., 2005), the turtle postnatal spinal cord (Reali et al., 2011) and the zebrafish spinal ECL (Chang et al., 2021). The GABAaR mediated excitation can induce increases in intracellular Ca2+ (Tozuka et al., 2005) (Reali et al., 2011) and influence proliferation and differentiation. We have shown that ECs in mammalian spinal cord depolarise to GABA through activation of GABAaRs, but with no GABAbR mediated responses (Corns et al., 2013). Since such depolarisations elicit calcium rises (Reali et al., 2011), there is potential for long lasting intracellular effects of GABAaR activation. Curiously, activation of nicotinic cholinergic receptors in mammalian ECs also elicits a depolarisation, yet results in increased proliferation (Corns et al., 2015). In zebrafish, acetylcholine acts on NSCs synaptically, whilst ambient GABA reduces their proliferation (Chang et al., 2021). However, the molecular pathways underpinning the differential outcomes on proliferation remain unclear and therefore require unravelling in future studies.

### 4.2 What is the source of GABA?

In the ECL, neighboring CSFcCs are GABAergic cells which possess the machinery for GABA synthesis and secretion (Corns et al., 2013; Gotts et al., 2016; Roberts et al., 1995; Tamamaki et al., 2003). Thus, CSFcCs may provide a source of GABA restricting proliferation of ECs, similarly to the role neuroblasts play in the SVZ (Daynac et al., 2013; Liu et al., 2005; Nguyen et al., 2003; Wang et al., 2003) where radiation induced death of fast dividing GABAergic progenitors activates dormant NSCs (Daynac et al., 2013). CSFcCs in lamprey have recently been suggested to be able to release somatostatin from dense core vesicles, although GABA release was not detected (Jalalvand et al., 2022). Future studies may be able to take advantage of mouse lines expressing cre recombinase in the CNS only in CSFcCs (Jurcic et al., 2021) to control their activity through opto- or chemo-genetics, to determine if they do indeed influence EC proliferation and/or differentiation.

### 4.3 DBI function in the spinal cord

In other neurogenic niches, the activity of GABA is modulated by the endozepine DBI, which acts to dampen the GABAergic ‘brake’ so the balance of activity in these two mechanisms provides sensitive control of overall neurogenic niche activity (Alfonso et al., 2012; Dumitru et al., 2017). Here we show that DBI and ODN are also expressed in the spinal cord. In the spinal cord both nestin+ and GFAP^+^ glia were DBI^+^, suggesting that astroglia in the spinal cord may also synthesize and release DBI, similar to that observed in the brain (Christian and Huguenard, 2013; Gandolfo et al., 2000; Loomis et al., 2010; Patte et al., 1999). We also show that the Sox2^+^ ECs of the CC region express high levels of both DBI and ODN similar to the SVZ and SGZ, which highlights the importance of endogenous DBI and its metabolites in the maintenance of adult NSC niches (Alfonso et al., 2012; Bormann, 1991; Tozuka et al., 2005).

Our data in flumazenil and Ro15-4513 treated animals or G2F77I mice with altered binding affinity of ligands to CBR show that proliferation is increased, while differentiation to glial cells is reduced when endogenous CBR ligand binding is modulated. This results in a larger population of undifferentiated Sox2^+^ NSC-like cells. These data reflect the role of DBI and ODN reported in the SVZ and SGZ (Alfonso et al., 2012; Dumitru et al., 2017). We now show that similar mechanisms involving endozepinergic modulation of adult proliferation are present in the spinal cord, but critically, we show that the mechanism of action of these endogenous proteins <u>may differ in the different regions</u>. Interruption or blocking of benzodiazepine binding site agonists to CBR, in G2F77I mutant- and flumazenil treated animals, respectively, reduces GABAergic tone on proliferation and thus proliferation increases. Furthermore, inverse agonist functionality requires constitutive receptor activity (Kenakin, 2001; Khilnani and Khilnani, 2011), therefore Ro15-4513 could be argued to reverse CBR mediated inhibitory GABAergic tone, increasing proliferation, and decreasing differentiation with a greater population of undifferentiated NSC-like Sox2+ cells.

Interestingly, in the SGZ and SVZ, DBI and ODN act as negative allosteric modulators (Alfonso et al., 2012; Dumitru et al., 2017) however, in the spinal cord the endozepinergic molecule is acting as a positive allosteric modulator. This difference in direction of effect of DBI is a well-known phenomenon in the reticular thalamic nucleus, where DBI is secreted by nearby astrocytes to positively influence GABAergic tone (Christian et al., 2013; Song et al., 2013). Glial cells of the spinal cord also express DBI and so a similar paracrine feedback loop may also exist within the spinal cord to regulate proliferation and differentiation.

### 4.4 The potential clinical significance of modulation of NSC proliferation

Our results show that decreasing the GABAergic tone on ECs results in an increase in their proliferation. The proliferative and lesion sealing response of ECs following spinal cord injury, and how these events can be manipulated for regeneration, has long been an exciting avenue of research (Moreno-Manzano, 2020). Although some work has shown that EC proliferation and astrocytic differentiation is inhibitory to spinal cord regeneration (Yang et al., 2020), others have shown poorer functional outcomes and greater tissue damage without the EC response (Sabelstrom et al., 2013). Our findings highlight the prospect of modulating EC proliferation with commonly available, clinically approved, GABA receptor modulators to examine effects on spinal cord function in disease and following injury.

Whilst GABAergic modulation can increase the number of cells in the spinal cord, further work is required to determine how best to direct differentiation for repair following cell depletion in times of injury. For example, in a mouse model of demyelination, reactive astrocytes were converted to myelinating mature oligodendrocytes by injection of Sox10 into the lesion site (Khanghahi et al., 2018; Shao et al., 2020). Recent work which integrates single-cell RNA sequencing (scRNA-seq) and single-cell assay for transposase-accessible chromatin using sequencing (scATAC-seq) has shown that ECs possess a latent genetic program for oligodendrogenesis which can be reactivated in mice genetically engineered to express OLIG2 in adult ECs. Following injury, OLIG2 expressing ECs exhibited enhanced oligodendrocytic differentiation, where these EC-derived oligodendroctyes were able to produce oligodendrocytic progeny themselves in addition to developing into mature myelinating oligodendrocytes, which contributed to the normalization of axon condition after SCI. Furthermore, ependymal oligodendrocyte generation occurred in parallel to the normal astrocytic differentiation seen in post-injury proliferative ECs which has been reported to be essential for lesion sealing to restrict tissue damage (Llorens-Bobadilla et al., 2020; Sabelstrom et al., 2013).

### 4.5 Can GABAergic signalling limit pathogenic proliferation in the spinal cord?

Controlling NSC quiescence is extremely important, as unchecked proliferation may lead to tumorigenesis and GABAergic signalling may hold the key to such regulation. Changes in GABAergic signalling, GABA receptor- and GABAaR subunit expression occur in various CNS tumours (Young and Bordey, 2009; Zhang et al., 2014; Kallay et al., 2019) where expression and activity of GABAaR negatively correlates with tumour grade (Smits et al., 2012) and is associated with attenuation of disease progression and improved survival (Blanchart et al., 2017). Furthermore, GABAergic modulators such as diazepam and midazolam inhibit proliferation of malignant glioma derived cell lines (Chen et al., 2013, 2016).

It has been hypothesized that endozepines may also be involved in the high proliferative rate of anaplastic tumour cells, as there is a significant increase in DBI expression in various brain tumours, including astrocytomas, glioblastomas, medulloblastomas, and ependymoma (Ferrarese et al., 1989; Alho, H, Kolmer, M, Harjuntausta, T, Helen, 1995; Miettinen et al., 1995; Duman et al., 2019). DBI also supports malignant glioma proliferation by modulating mitochondrial long-chain fatty acyl-CoA availability which promotes fatty acid β-oxidation mediated tumour growth (Duman et al., 2019). Whilst the effects of DBI upon GABAergic signalling are clear in physiological proliferation of NSCs, further work is required to determine whether this pathway is also relevant in pathophysiological proliferation. DBI overexpression in glioma could contribute to inhibition of GABAergic signalling in low-grade tumour cells to promote tumour growth (Jung et al., 2019) where modulation of endogenous CBR site binding by antagonists such as flumazenil could be investigated as a means to reduce aberrant proliferation.

### 4.6 Summary

In conclusion, proliferation in the adult spinal cord may be modulated by GABAergic signaling and modulation by endozepines such as DBI may be one of the mechanisms restricting proliferation of cells in the neurogenic niche surrounding the CC of the mammalian spinal cord. NSC properties of ECs should not be overlooked when considering the possibility of self-regenerative therapies in future clinically relevant studies for spinal cord repair, and that these cells, and the niche signaling which influences the proliferative capacity of the intact cord warrants further investigation.

## Acknowledgements

Tissue from G2F77I animals was kindly provided by Dr I Dumitru and Professor H. Monyer (DFKZ Heidelberg, Germany)

Research involving confocal microscopy was kindly supported by The Wellcome Trust WT104918MA

Lauryn New was supported by a University Research Scholarship

This publication uses data collected within the framework of the PhD thesis GABAergic regulation of proliferation in the postnatal spinal cord of Lauryn New published in 2020 at the Faculty of Biological Sciences, University of Leeds, UK

## Abbreviations

CBR: central benzodiazepine receptor
CC: central canal
CSFcCs: cerebrospinal fluid contacting cells
DAPI: 4’,6-diamidino-2-phenylindole
DBI: Diazepam binding inhibitor
DMSO: Dimethyl sulfoxide
EDTA: ethylenediamine tetraacetic acid
EdU: Ethynyl-2’-deoxyuridine
EC: Ependymal cells
ECL: ependymal cell layer
GABA: γ-aminobutyric acid
GABAaR: GABAa receptor
GAD67-GFP: glutamic acid decarboxylase67-green fluorescent protein
GFAP: glial fibrillary acidic protein
GM: grey matter
HPLC: high performance liquid chromatography
IP: intraperitoneal
IR: immunoreactivity
NSC: neural stem cell
ODN: octadecaneuropeptide
PFA: paraformaldehyde
PCA: perchloric acid
PB: phosphate buffer
PBST: PBS containing 0.1% triton
SGZ: subgranular zone
SVZ: subventricular zone
VGB: vigabatrin
WM: white matter
WT: wild type

